# Biophysical characterizations of the recognition of the AAUAAA polyadenylation signal

**DOI:** 10.1101/503755

**Authors:** Keith Hamilton, Yadong Sun, Liang Tong

## Abstract

Most eukaryotic messenger RNA precursors must undergo 3’-end cleavage and polyadenylation for maturation. We and others recently reported the structure of the AAUAAA polyadenylation signal (PAS) in complex with the protein factors CPSF-30, WDR33 and CPSF-160, revealing the molecular mechanism for this recognition. Here we have characterized in detail the interactions between the PAS RNA and the protein factors using fluorescence polarization experiments. Our studies show that AAUAAA is recognized with ~1 nM affinity by the CPSF-160–WDR33–CPSF-30 ternary complex. Variations in the RNA sequence can greatly reduce the affinity. Similarly, mutations of residues that have van der Waals interactions with the bases of AAUAAA also lead to substantial reductions in affinity. Finally, our studies confirm that both CPSF-30 and WDR33 are required for binding the PAS RNA, and determine a ~7 nM affinity between CPSF-30 and the CPSF-160–WDR33 binary complex.

## Introduction

Most eukaryotic messenger RNA precursors (pre-mRNAs) must undergo extensive processing to become functional mRNAs, which includes 5’-end capping, splicing, and 3’-end cleavage and polyadenylation (1–5). The recognition of a polyadenylation signal (PAS) is a crucial step for 3’-end processing, which helps to define the position of cleavage in the pre-mRNA as the PAS is often located 10-30 nucleotides upstream of the cleavage site. The PAS is a hexanucleotide, and the most common motif is AAUAAA (~55% frequency) for mammalian pre-mRNAs, followed by the AUUAAA motif (~16% frequency) (6–8). Many other motifs can also support 3’-end processing, but are much rarer (<4% frequency). AAUAAA, AUUAAA and 10 other single nucleotide variants account for ~92% of PAS in human and mouse pre-mRNAs (8).

Many proteins are involved in pre-mRNA 3’-end processing (9–11), and several subcomplexes of this 3’-end processing machinery have been identified, including the cleavage and polyadenylation specificity factor (CPSF) and the cleavage stimulation factor (CstF). The 73-kDa subunit of CPSF (CPSF-73) is the endoribonuclease for the cleavage reaction (12), and two other CPSF subunits, WDR33 (11) and CPSF-30, are required for recognizing the PAS (13,14). CPSF-30 also interacts with Fip1, another subunit of CPSF, which helps to recruit the poly(A) polymerase to the 3’-end processing machinery. CstF recognizes a G/U-rich sequence motif downstream of the cleavage site, and it also has an important role in alternative polyadenylation (15–17).

We and others recently reported the structures of a quaternary complex of human CPSF-160, CPSF-30, WDR33 and an AAUAAA PAS RNA (Figs. 1A, 1B) (18,19), the structure of a ternary complex of the yeast protein homologs (Cft1, Yth1 and Pfs2, without RNA) (20), as well as the structure of a binary complex of human CPSF-160 and WDR33 (21). The structures of the quaternary complexes revealed extensive and specific interactions between the AAUAAA PAS and WDR33 and CPSF-30 (Fig. 1C), while CPSF-160 serves a crucial scaffolding role in the complex. In addition, there is a Hoogsteen base pair between U3 and A6 of the PAS. A few aspects of the interactions between the ternary complex and PAS RNA have been studied by fluorescence polarization assays (21).

**Figure 1.**
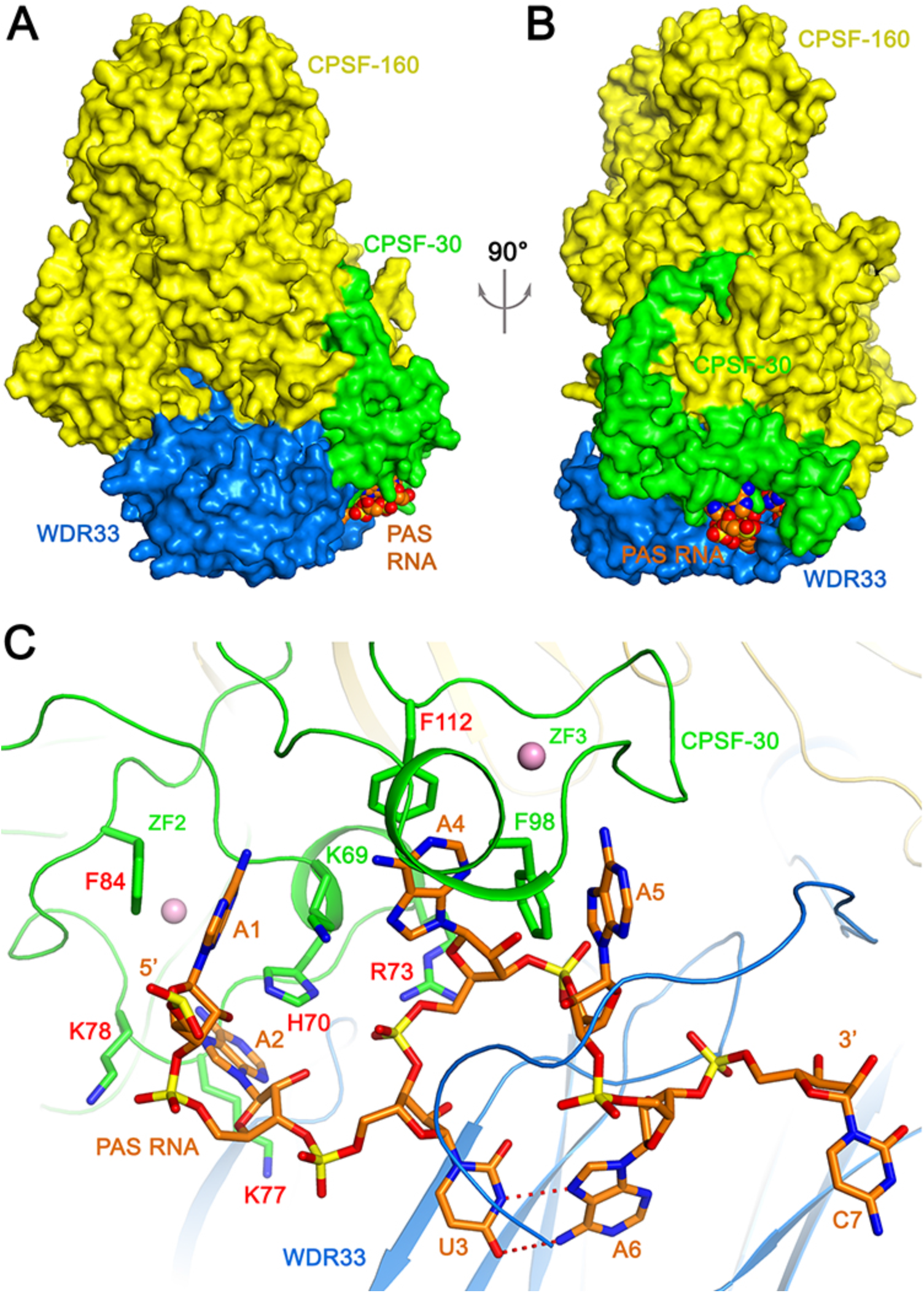
Overall structure of the human CPSF-160–WDR33–CPSF-30–PAS RNA quaternary complex. **(A)**. Schematic drawing of the quaternary complex. CPSF-160 (yellow), WDR33 (blue) and CPSF-30 (green) are shown as molecular surfaces. The PAS RNA is shown as a sphere model (orange). **(B)**. Structure of the quaternary complex, viewed after a 90° rotation of around the vertical axis. **(C)**. Recognition of the AAUAAA PAS (orange) by CPSF-30 zinc fingers ZF2-ZF3 (green) and WDR33 (blue). Hydrogen-bonds in the U3-A6 Hoogsteen base pair are indicated with dashed lines in red. Side chains of CPSF-30 that contact the RNA bases are shown as stick models, and those selected for mutagenesis studies are labeled in red. Zinc atoms are shown as spheres in pink. Produced with PyMOL (www.pymol.org).

We report here detailed characterizations of the interactions between CPSF-160, WDR33, CPSF-30, and various PAS RNAs. We show that the ternary complex has high affinity for the AAUAAA PAS RNA, with *K*_d_ of ~1 nM, and the AUUAAA PAS RNA has a *K*_d_ of ~10 nM. In comparison, other sequence motifs that can also support 3’-end processing, as well as changes to the U3-A6 Hoogsteen base pair, lead to substantial reduction in the binding affinity. In addition, mutations of CPSF-30 residues that are in contact with the RNA bases can also give rise to reductions in binding affinity. The CPSF-160–WDR33 and CPSF-160–CPSF-30 binary complexes have much lower affinity for the RNA, confirming that both WDR33 and CPSF-30 are required for PAS recognition. The CPSF-160–WDR33 binary complex has high affinity for CPSF-30, with a *K*_d_ of ~7 nM.

## Results

### Variation of the AAUAAA RNA length

The structures of the quaternary complexes show that WDR33 and CPSF-30 primarily recognize the PAS hexanucleotide itself (Fig. 1C). Although a 17-mer RNA was used for the structural study, only the PAS was found to be well ordered (18). The nucleotide directly following the PAS was weakly ordered, and the other nucleotides were disordered. To assess this structural observation, we used RNA oligos of various lengths, 17-mer (FAM-AACCUCC<u>AAUAAA</u>CAAC), 11-mer (FAM-CCUCC<u>AAUAAA</u>CA) and 6-mer (<u>AAUAAA-FAM</u>), and carried out fluorescence polarization binding assays. All these oligo RNAs carry a FAM fluorescent label at the 5’ or 3’ end, allowing direct observation of the fluorescence polarization signal.

The experimental data confirmed that the 17-mer and 11-mer RNAs have nearly the same binding affinity to the CPSF-160–WDR33–CPSF-30 ternary complex, with *K*_d_ values of 0.56 nM and 1.4 nM, respectively (Fig. 2A). The 17-mer RNA was used in titration experiments numerous times, and the observed *K*_d_ values ranged between 0.6 and 1.2 nM, consistent with that reported in an earlier study (21). Fip1 (residues 137-243) was included at 10 μM concentration in all the assays to stabilize the C-terminal segment of CPSF-30, although it did not have an effect on RNA binding.

**Figure 2.**
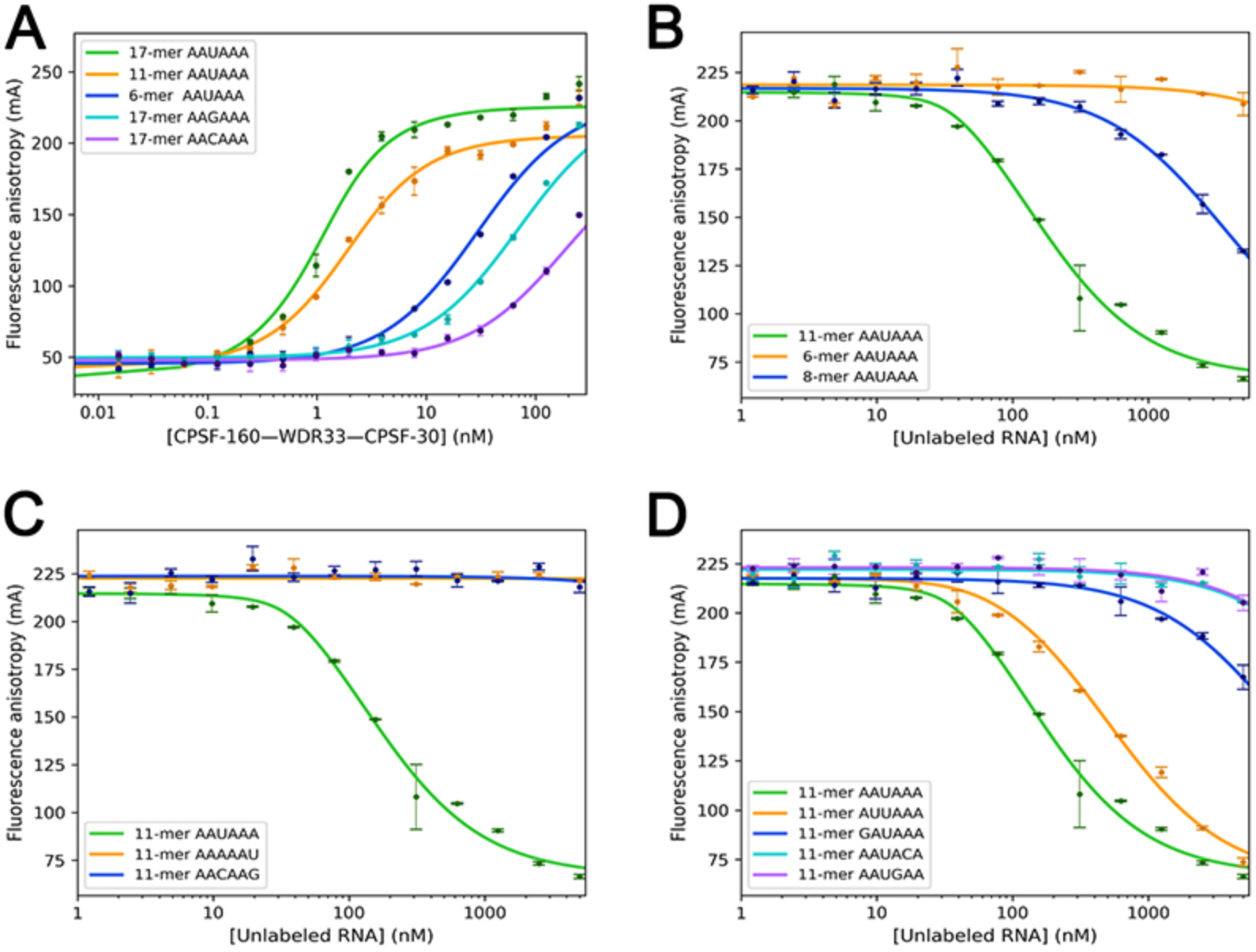
Effects of variations in the RNA on PAS recognition. **(A)**. Fluorescence polarization binding assays of the CPSF-160–WDR33–CPSF-30 ternary complex with labeled 17-mer, 11-mer and 6-mer AAUAAA PAS RNAs, as well as labeled 17-mer RNAs with AA<u>G</u>AAA and AA<u>C</u>AAA as the PAS (nucleotides distinct from AAUAAA are indicated by underline). The curves represent theoretical fitting to the binding data. **(B)**. Competition fluorescence polarization binding assays of the CPSF-160–WDR33–CPSF-30 ternary complex with unlabeled 11-mer, 8-mer and 6-mer AAUAAA PAS RNAs. Labeled 17-mer AAUAAA PAS RNA was used as the reporter. **(C)**. Competition fluorescence polarization binding assays of the CPSF-160–WDR33–CPSF-30 ternary complex with unlabeled 11-mer RNAs containing variations of the U3-A6 Hoogsteen base pair, AA<u>A</u>AA<u>U</u> and AA<u>C</u>AA<u>G</u>. **(D)**. Competition fluorescence polarization binding assays of the CPSF-160–WDR33–CPSF-30 ternary complex with unlabeled 11-mer RNAs containing variations of the A1, A2, A4 and A5 bases. Error bars are standard deviations from two repeats.

In comparison, the 6-mer oligo had a higher *K*_d_ value of 29 nM. This could be due to interference by the FAM label or contribution from nucleotides outside the PAS. For example, the phosphate group of the nucleotide directly following the PAS has some interactions with WDR33 (Fig. 1C). To assess these different scenarios, we used an unlabeled AAUAAA 6-mer oligo and an unlabeled CAAUAAAC 8-mer oligo and carried out competition fluorescence polarization binding assays against the 17-mer FAM-labeled oligo (Fig. 2B). The *K*_d_ value determined from this assay for the AAUAAA 8-mer oligo was 72 nM, 24-fold higher than the 11-mer oligo, while the 6-mer oligo showed some binding only at 5000 nM concentration of the ternary complex. These data indicate that nucleotides outside the AAUAAA hexamer also contribute significantly to the binding.

The phosphate group directly following the PAS is likely a major factor enhancing the binding affinity. The FAM label in the 6-mer oligo is located at the 3’-end, which contains a phosphate group at that position, consistent with the higher affinity of this oligo (Fig. 2A). We carried out the competition assay with the unlabeled 11-mer oligo (Fig. 2B), and obtained a *K*_d_ value of 3.0 nM, indicating that the FAM label did not have a significant effect on the binding affinity of this longer RNA (the *K*_d_ for the FAM-labeled 11-mer RNA was 1.4 nM, Fig. 2A). Overall, these assays demonstrate that AAUAAA plus the phosphate group of the following nucleotide is crucial for high-affinity binding. Other nucleotides make smaller contributions to the interaction.

### Variation of the U3-A6 Hoogsteen base pair

A U3-A6 Hoogsteen base pair was observed in the PAS when bound to the CPSF-160–WDR33–CPSF-30 ternary complex (Fig. 1C) (18,19). The binding mode of the PAS suggests that other Hoogsteen base pairs, such as C3-G6, could be accommodated, while a wobble U3-G6 base pair would not fit. To assess whether the alternative Hoogsteen base pair can support binding to the ternary complex, we determined the binding affinity of unlabeled 11-mer RNA with AA<u>C</u>AA<u>G</u>(variations from the AAUAAA PAS are indicated by underlines) as the equivalent of the AAUAAA PAS by competition fluorescence polarization assays. The experimental data showed no binding of the oligo even at 5000 nM concentration of the ternary complex (Fig. 2C), indicating that the alternative Hoogsteen base pair could not be accommodated in the binding site. The 11-mer RNA with AA<u>A</u>AA<u>U</u> as the PAS showed no binding either.

We also characterized the binding of FAM-labeled 17-mer RNAs with AA<u>G</u>AAA and AA<u>C</u>AAA as the PAS, to test the effect of breaking the U3-A6 Hoogsteen base pair. These RNAs did show binding to the ternary complex, with *K*_d_ values of 69 and 180 nM, respectively (Fig. 2A), roughly 120- and 320-fold higher than AAUAAA. The *K*_d_ value for the AA<u>G</u>AAA oligo is consistent with that reported in an earlier study (21).

### Variation of other positions of the AAUAAA PAS

Besides the U3-A6 Hoogsteen base pair, the structures show that A1 and A4 of the AAUAAA PAS are specifically recognized by CPSF-30, while A2 and A5 also establish favorable hydrogen-bonding interactions (18,19). To assess the binding affinity of other PAS hexamers that can also support 3’-end processing, we selected from those identified in mammalian pre-mRNAs (7,8), changing each of these four positions at a time. The unlabeled 11-mer RNAs that we studied included <u>G</u>AUAAA (~1% frequency, first position), A<u>U</u>UAAA (~16%, second position), AAU<u>G</u>AA (~1%, fourth position), and AAUA<u>C</u>A (~2%, fifth position) as the PAS.

The experimental data showed that these variant PAS RNAs have much lower affinity for the ternary complex, except for A<u>U</u>UAAA, which is the second most frequently observed PAS (Fig. 2D). The *K*_d_ value for A<u>U</u>UAAA 11-mer oligo is 10.3 nM, only about 3-fold higher than the corresponding 11-mer AAUAAA oligo (*K*_d_ of 3.0 nM). The <u>G</u>AUAAA 11-mer oligo has a *K*_d_ value of 170 nM (55-fold higher), while the AAU<u>G</u>AA and AAUA<u>C</u>A 11-mer oligos showed only minor binding at 5000 nM concentration (Fig. 2D). AA<u>G</u>AAA is another PAS hexamer with low frequency (~3%), and it had *K*_d_ of 69 nM (120-fold higher) (Fig. 2A).

### Mutations of CPSF-30

Besides hydrogen-bonding interactions between the adenine bases and the backbone of CPSF-30, there are also extensive van der Waals interactions. Specifically, the A1, A2, A4 and A5 bases are each involved in π-stacking interaction with an aromatic side chain of CPSF-30: A1 with Phe84, A2 with His70, A4 with Phe112, and A5 with Phe98 (Fig. 1C). In addition, A2 is flanked on the other face by two Lys side chains, Lys77 and Lys78, and Arg73 has ionic interactions with the phosphate connecting nucleotides U3 and A4.

To test the importance of these interactions for PAS RNA binding, we produced the H70A, R73A, K77A/K78A, F84A and F112A mutants of CPSF-30. Each mutant and the wild-type CPSF-30 was mixed with the CPSF-160–WDR33 binary complex at 10-fold molar excess, which we showed was sufficient to achieve nearly maximal binding to the RNA (see next). The affinity of these mixtures for the FAM-labeled 17-mer RNA was then determined (Fig. 3). The K77A/K78A mutant had roughly 3-fold higher *K*_d_ (2.8 nM) compared to wild-type CPSF-30. On the other hand, the F84A and R73A mutations had larger effects on the binding, with 35- and 55-fold higher *K*_d_ values (32 and 49 nM). Finally, the H70A and F112A mutants showed the largest effects, with 440- and 590-fold higher *K*_d_ values (390 and 530 nM).

**Figure 3.**
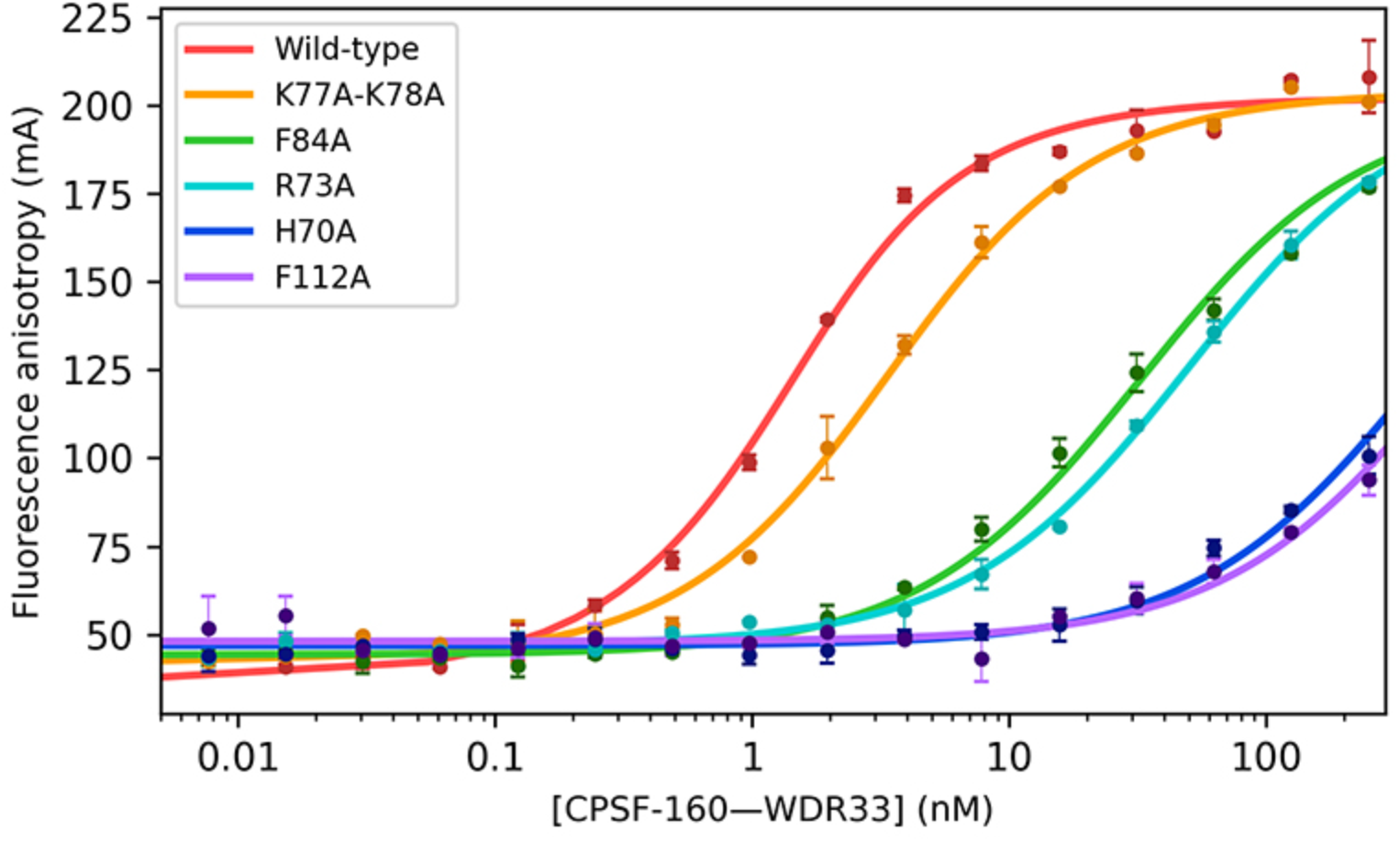
Effects of mutations in CPSF-30 on PAS recognition. Fluorescence polarization binding assays between the labeled 17-mer AAUAAA PAS RNA and mixtures of the CPSF-160–WDR33 binary complex and CPSF-30 wild-type and mutants at 10-fold molar ratio. Error bars are standard deviations from two repeats.

### Requirement of both WDR33 and CPSF-30 for RNA binding

The AAUAAA RNA is bound at the interface between WDR33 and CPSF-30 (Fig. 1C), and our earlier electrophoretic mobility shift assays showed that CPSF-30 alone, or the CPSF-160–WDR33 and CPSF-160–CPSF-30 binary complexes could not bind the RNA (18). To characterize these interactions more quantitatively, we mixed CPSF-30 at increasing molar ratios (0-, 0.5-, 1-, 2-, 5-, 10-, 20- and 30-fold) with the CPSF-160– WDR33 binary complex, and observed a clear enhancement of the apparent affinity of the mixture for the RNA when CPSF-30 concentration was increased (Fig. 4). Above 10-fold molar ratio of CPSF-30 relative to CPSF-160–WDR33, nearly maximal RNA binding was obtained. On the other hand, in the absence of CPSF-30, no binding was observed even at 125 nM concentration of the CPSF-160–WDR33 binary complex. Similarly, no RNA binding was observed for CPSF-30 alone at up to 125 nM concentration, consistent with both WDR33 and CPSF-30 being required for RNA binding.

**Figure 4.**
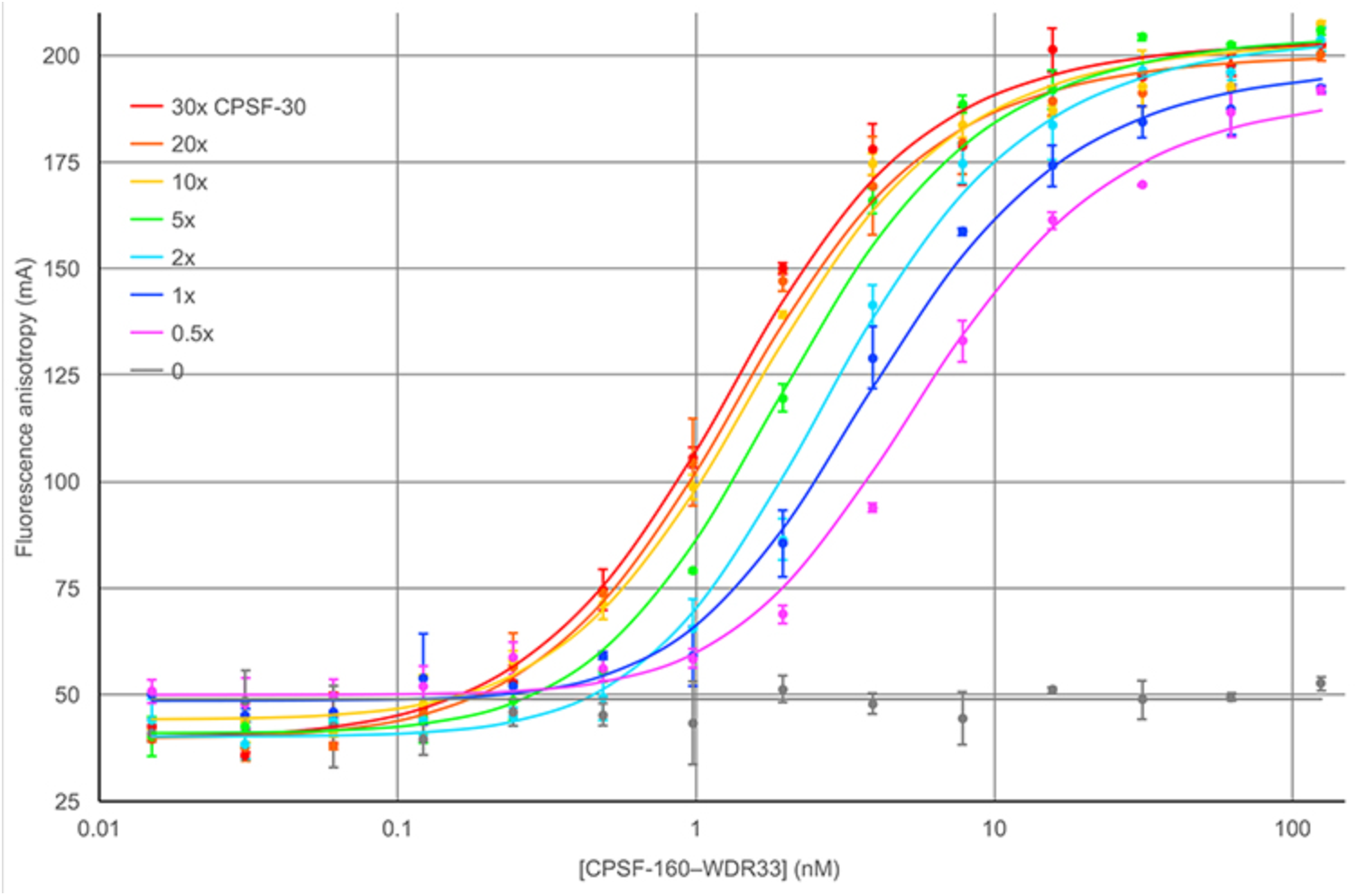
Both CPSF-30 and WDR33 are required for PAS RNA binding. Fluorescence polarization binding assays between the labeled 17-mer AAUAAA PAS RNA and mixtures of the CPSF-160–WDR33 binary complex and CPSF-30 at increasing molar ratios. Error bars are standard deviations from two repeats.

Assuming that each mixture contains two equilibria: A+B=AB and AB+C=ABC, where A represents the CPSF-160–WDR33 binary complex, B represents CPSF-30, and C represents the RNA, the total concentrations of CPSF-160–WDR33, CPSF-30 and RNA in the mixture and the *K*_d_ values for the two equilibria would determine the shape of each titration curve, which varied the concentration of CPSF-160–WDR33 (A) while keeping the molar ratio of CPSF-30 (B) constant. The *K*_d_ of the ternary complex for the RNA (ABC) was fixed at 0.6 nM based on the titration data for this complex (Fig. 2A). We carried out a global fit of all the titration curves, which determined the *K*_d_ for CPSF-30 binding to CPSF-160–WDR33 as 7.1 nM (Fig. 4).

## Discussion

Overall, our studies have provided detailed knowledge on PAS recognition by the CPSF-160–WDR33–CPSF-30 ternary complex, extending beyond the binding study reported earlier (21). The results confirm that both hydrogen-bonding and van der Waals interactions are important for the recognition of the A1, A2, A4 and A5 bases. Changing the identity of these bases generally has strong deleterious effects on the binding affinity, with the exception of the AUUAAA sequence, consistent with it also being frequently observed for 3’-end processing. On the other hand, loss of π-stacking interactions with these bases is also detrimental for the recognition. The structure suggests that the U2 base of AUUAAA could maintain the hydrogen-bond with the main-chain amide of Lys78 in CPSF-30 with a small conformational change, as well as the π-stacking with His70 (Fig. 1C).

Our studies demonstrate the importance of the U3-A6 Hoogsteen base pair for high-affinity binding to the ternary complex. The AA<u>A</u>AA<u>U</u> hexamer, swapping the positions of the U3 and A6 nucleotides, cannot maintain a Hoogsteen base pair as the AAUAAA hexamer, explaining the lack of binding for this RNA. On the other hand, the AA<u>C</u>AA<u>G</u> hexamer appears to fit nicely into the binding site, with the guanine base flanked on either side by Phe43 and Phe153 of WDR33 and picking up a hydrogen-bond between its 2-amino group and the main-chain carbonyl of Thr115 in WDR33. The exact reason why this hexamer cannot bind with high affinity is not clear, although it is consistent with the fact that it is not frequently observed for 3’-end processing (8). At the same time, the AA<u>G</u>AAA and AA<u>C</u>AAA hexamers are able to bind the ternary complex, albeit with substantially reduced affinity, indicating that a base pair here may not be absolutely required for binding.

The less frequently observed PAS motifs studied here appear to have much lower affinity for the ternary complex. In fact, the AAUAAA hexamer is most often associated with the last PAS in human and mouse pre-mRNAs, while the less frequently observed hexamers are associated with upstream PAS of pre-mRNAs with two or more processing sites (8). Especially, the AA<u>G</u>AAA hexamer is often found as the PAS in an upstream exon. This suggests that these less frequently observed PAS hexamers have a more prominent role in alternative polyadenylation, and that other protein factors (such as CstF) as well as recognition of auxiliary sequence motifs may be important for the processing at these sites.

## Methods

### Protein expression and purification

A human CPSF-160–WDR33 binary complex was expressed in baculovirus-infected Hi5 insect cells and a full-length human CPSF-30 was expressed in *E. coli* as an MBP fusion protein as described earlier (18). The components were mixed together for the binding assays, allowing the variation of the molar ratios of CPSF-30 relative to CPSF-160–WDR33.

A CPSF-160–WDR33–CPSF-30 ternary complex was expressed in Hi5 insect cells with the Multibac expression system (22) (Geneva Biotech). WDR33 (residues 1-425) carried an N-terminal His-tag, followed by MBP and a TEV protease cleavage site. CPSF-160 and CPSF-30 are untagged.

Human Fip1 (residues 137-243) was cloned into the pET28a vector (Novagen), with N-terminal His and SUMO tags, and expressed in *E. coli* BL21(DE3) cells.

The CPSF-160–WDR33–CPSF-30 and the CPSF-160–WDR33 complexes were purified following the same protocol. The insect cells were lysed by sonication in a buffer containing 50 mM Tris (pH 8.0), 500 mM NaCl, 30 mM imidazole, 10 mM beta-mercaptoethanol, and one SIGMAFAST protease inhibitor cocktail tablet. The lysate was mixed with 2 mL Ni-NTA beads (Qiagen), washed with 15 mL buffer containing 2 M NaCl to remove bound nucleic acids, then the protein was eluted with 5 mL buffer containing 250 mM imidazole. 1 mg TEV protease was added, and the sample was incubated overnight at 4°C. It was then run over a Superdex200 16/60 column (GE Healthcare), using a buffer containing 20 mM (Tris 8.0), 350 mM NaCl, and 10 mM DTT.

For Fip1, the cells were lysed by sonication in a buffer containing 50mM Tris (pH 8.0), 200 mM NaCl, 30 mM imidazole, 10 mM beta-mercaptoethanol, and 2 mM PMSF. The lysate was incubated with 5 mL Ni-NTA, washed with 30 mL buffer, then Fip1 was eluted with 10 mL buffer containing 250 mM imidazole. 100 μg UlpI protease was added, and the sample was incubated at 4°C for one hour. It was then purified further using a 5 mL Fastflow MonoQ column followed by a Superdex 200 16/60 column using a buffer containing 20 mM Tris (pH 8.0), 100 mM NaCl, and 10 mM DTT.

### Fluorescence polarization binding assays

The assays were performed using a Neo2S plate reader (Biotek). The buffer for all the assays contained 20 mM Tris (pH 8.0), 150 mM NaCl, 10 mM DTT, and 10 μM Fip1. A 17-mer oligonucleotide that included a polyadenylation site and the surrounding bases from the SV40 virus and a 6-carboxyfluorescein (6-FAM) 5’-end label was used as the probe at a concentration of 1 nM in all experiments. For assays that involved titration with CPSF-30, various molar ratios of CPSF-30 were added to CPSF-160–WDR33 and the mixture was allowed to incubate on ice for 1 h. Oligonucleotides were purchases from IDT. The data from competition experiments were fitted using an analytical equation (23).

## Acknowledgments

This research is supported by NIH grant R35GM118093 (to LT).

